# Sequence Coverage Visualizer: A web application for protein sequence coverage 3D visualization

**DOI:** 10.1101/2022.01.12.476109

**Authors:** Xinhao Shao, Christopher Grams, Yu Gao

**Affiliations:** Department of Pharmaceutical Sciences, University of Illinois at Chicago; Department of Computer Sciences, University of Illinois at Chicago

**Keywords:** 3D structure, protein sequence coverage, visualization, proteomics

## Abstract

Protein structure is connected with its function and interaction and plays an extremely important role in protein characterization. As one of the most important analytical methods for protein characterization, Proteomics is widely used to determine protein composition, quantitation, interaction, and even structures. However, due to the gap between identified proteins by proteomics and available 3D structures, it was very challenging, if not impossible, to visualize proteomics results in 3D and further explore the structural aspects of proteomics experiments. Recently, two groups of researchers from DeepMind and Baker lab have independently published protein structure prediction tools that can help us obtain predicted protein structures for the whole human proteome. Although there is still debate on the validity of some of the predicted structures, it is no doubt that these represent the most accurate predictions to date. More importantly, this enabled us to visualize the majority of human proteins for the first time. To help other researchers best utilize these protein structure predictions, we present the Sequence Coverage Visualizer (SCV), http://scv.lab.gy, a web application for protein sequence coverage 3D visualization. Here we showed a few possible usages of the SCV, including the labeling of post-translational modifications and isotope labeling experiments. These results highlight the usefulness of such 3D visualization for proteomics experiments and how SCV can turn a regular result list into structural insights. Furthermore, when used together with limited proteolysis, we demonstrated that SCV can help validate and compare different protein structures, including predicted ones and existing PDB entries. By performing limited proteolysis on native proteins at various time points, SCV can visualize the progress of the digestion. This time-series data further allowed us to compare the predicted structure and existing PDB entries. Although not deterministic, these comparisons could be used to refine current predictions further and represent an important step towards a complete and correct protein structure database. Overall, SCV is a convenient and powerful tool for visualizing proteomics results.

## Introduction

To date, most of the proteomics experiments are conducted using the bottom-up proteomics method. Proteins are digested by sequence grade trypsin into smaller peptides and then separated and identified by LC-MS/MS. Comparing to other proteomics methods, bottom-up proteomics is by far the most efficient way to identify and quantify thousands of proteins within hours. However, since the analyte is not the whole protein but rather digested tryptic peptides, bioinformatics is needed to assemble all the available information to infer the original proteins. This entire process makes the protein sequence coverage, i.e., the ratio of total identified length to total protein length, an essential parameter for bottom-up proteomics. The average protein sequence coverage of a typical bottom-up proteomics experiment varies from 10% to 95%, depending on the complexity of the sample and the sample processing method. The coverage number generally increases monotonically as the digestion is more and more complete through time.

Noting this phenomenon, limited proteolysis methods have been developed to use various digestion time points to differentiate the easy-to-digest and hard-to-digest parts of the proteins1–3. It is reasonable to believe that the initial digested peptides are from the more accessible regions of the protein, and the late digested peptides are generally from hard-to-access regions. When a time series is created using limited proteolysis, we can easily map all the identified peptides from different time points to the protein sequence. Even better, if we could map all these peptides onto the 3D model of the protein, we could see the sequence coverage in 3D and validate if the identified peptides are from the same region or not. If the sequence coverage increases over time, we can further visualize it in space to see how trypsin digests the protein step by step. Such visualization will significantly improve our understanding of the protein structure and even help us to refine protein structures further. However, this was not entirely possible in the past, as crystallography, NMR, and Cryo-EM protein structures cover only a limited portion of the known proteome. In 2021, two groups of researchers, one from DeepMind and one from the Baker lab, published two powerful tools for protein structure prediction4–6. Although there is still debate on the validity of some of the predicted structures, it is no doubt that these represent the most accurate predictions we can get to date. More importantly, these tools and the published data enabled us to visualize the majority of human proteins for the first time.

These predictions enabled us to create this Sequence Coverage 3D Visualizer (SCV) as a free web application, to assist researchers worldwide in visualizing proteomics results in 3D. To demonstrate the usage and usefulness of SCV, we used multiple examples to highlight the features of SCV. Using previously published isotope labeling data, we showed that SCV can be used to visualize and compare differentially labeled isotopes. When used together with limited proteolysis of native proteins, SCV could visualize the digestion progress and therefore compare predicted structures to existing PDB entries. Such comparison may serve as an important step towards validating predicted structures from both Alphafold2 and RoseTTAFold. Overall, we demonstrated that SCV can be easily used to convert proteomics results into structural insights.

## Results

The Sequence Coverage 3D Visualizer (SCV) can be accessed at http://scv.lab.gy/. The basic function of SCV is to visualize protein sequence coverage in 3D using the peptide list identified from experiments (**Fig. 1**). SCV has a user-friendly web interface (**Fig. 1**) which allows the user to directly paste an experimentally identified peptide list. When post-translational modifications (PTMs) are included in the results as numbers in brackets (“[]”), they will be automatically detected and visualized with different colors. The background color and the color of each PTM can be customized easily through the interface. When a species is selected (currently supports human, mouse and rat data), all the peptides are mapped to the corresponding UniProt reference database. All detected proteins will be listed with gene name, UniProt identifier and a brief description of the protein. In addition, a linear sequence coverage map is generated for each protein entry to show the positions of detected peptides. When selected from the protein list, a rotating 3D structure is generated, with red color highlighting the covered sequence and other user-selected colors to highlight the PTMs. By default, SCV uses Alphafold’s predicted 3D models of the protein as it covers the majority of human proteome and provide good accuracy for most known proteins we tested. Custom 3D model mapping is supported through PDB file upload. When users upload a PDB file of a protein or protein complexes, the uploaded structure will be used for visualization instead of the default predicted structure.

**Figure 1.**
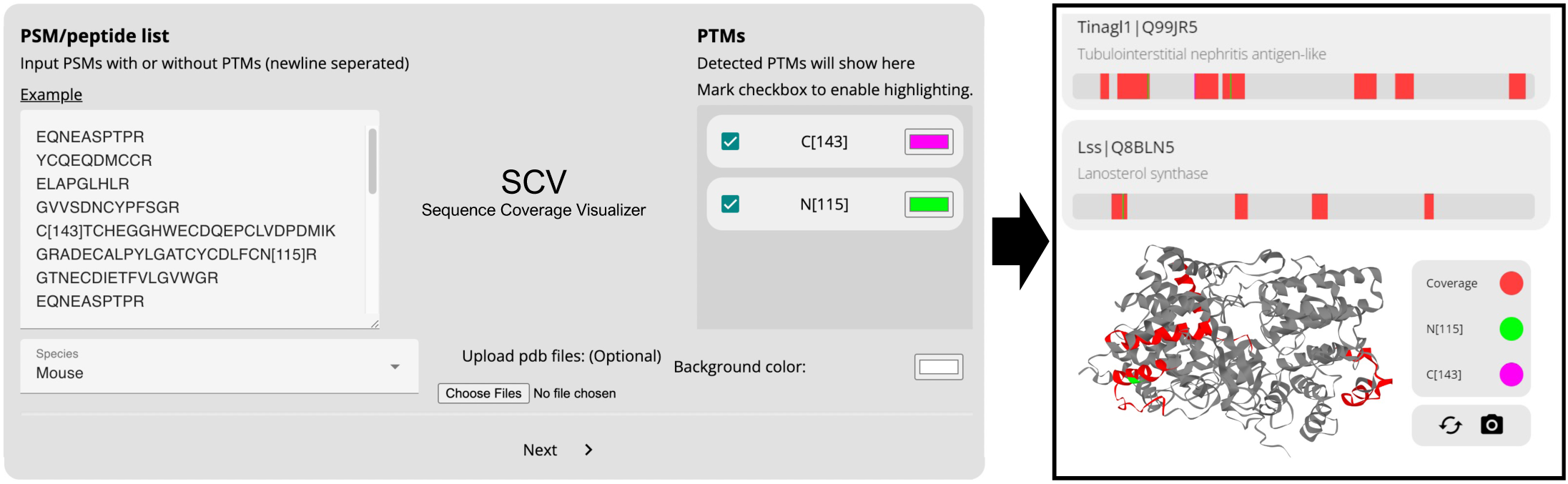
Overview of SCV web application input and output. SCV uses peptide list and species information to generate a list of proteins with linear sequence coverage and let users to select the protein to visualize in 3D. By default, Alphafold predicted structure is used; users can also upload PDB files to customize 3D model. Post-translational modifications are automatically detected and can also be visualized by custom colors.

Once submitted, the peptide list is then processed by our backend Node.js server and custom Python scripts running on our server. A typical response of a few thousand peptides and a few hundred proteins takes only seconds to process. A permanent link is provided to reference the result and it can be shared with others and inspected later. The protein list is sorted by sequence coverage percentage, from high to low. The 3D visualization is rendered using GLmol^7^, a WebGL library, and Three.js^8^, a JavaScript library for 3D graph display. Our interactive 3D visualization allows the user to move and rotate the protein to make close observations easily. SCV works on all major web browsers on both desktop and mobile devices.

### Isotope labeling visualization

SCV could be used to visualize and validate isotope-labeled proteomics results by inspecting the spatial relationship between differentially labeled residues. Here, we used the covalent protein painting data from Bamberger et al. ^9^ to demonstrate its usage. In this paper, Bamberger et al. described a method to differentially label “heavy” and “light” dimethyl groups to proteins before and after protein unfolding. In native form, proteins are first labeled in vivo with “light” dimethyl groups through reductive amination. The surface or more exposed lysine groups of the protein are therefore labeled. Later, with denaturing conditions, unfolded proteins are labeled again with “heavy” dimethyl groups. Because the more accessible lysine groups have already been labeled in the previous step, only the newly exposed lysine groups are labeled with the “heavy” dimethyl groups, making it possible to differentiate lysines with different accessibility. Here, we picked one of the proteins from the experiment, the GAPDH, as an example to show how SCV could be used to visualize different isotopes of the “heavy” and “light” dimethyl groups on lysine (**Fig. 2**). As shown in **Fig. 2a**, the whole GAPDH protein is visualized in 3D and the detected sequences are covered in red color. The green residues are the lysine groups labeled with the “light” label and the blue residues are the lysine groups labeled in “heavy”. This visualization result shows that mild heat shock will increase the accessibility of the lysine groups, as described in the paper. The interactive interface that allows free rotation and zoom in/out enabled us to inspect the result in space and better understand the spatial relationship between the native and unfolded forms of the protein. With the help of SCV, 3D visualization of both “heavy” and “light” labeling of all the protein described in this paper can be generated within seconds without any expert knowledge or the use of additional tools.

**Figure 2.**
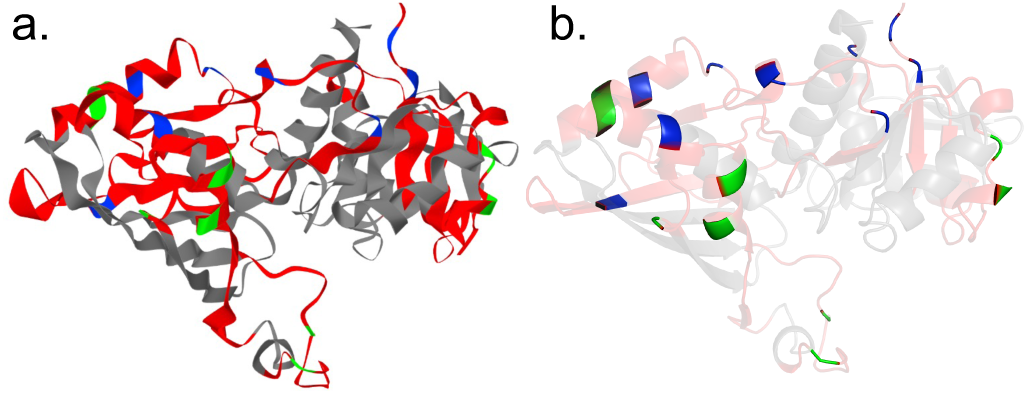
Isotope labeling visualization of GAPDH protein labeled with both heavy and light dimethyl. a) 3D visualization generated by SCV. Red: covered sequence; grey: uncovered sequence; green: light label; blue: heavy label. b) transparency adjusted model to highlight the position of the labeled residues.

### 3D visualization of digestion through time

Limited proteolysis has been used to obtain protein structural information in previous research. When a protein is only allowed to be digested by a protease for a limited time, easy-to-digest regions are detected in early time points while hard-to-digest regions are detected later. When plotted through time, the appearance of different peptides from different regions of the protein can be correlated to the accessibility of the region (**Fig. 3**). When proteins are digested in their native form, limited proteolysis can be used to determine the more accessible surfaces, which are often digested first, followed by the less accessible regions, till complete unfolding. Here, we performed native digestion of HEK293 cell lysate using trypsin at four different time points: 1, 2, 4, 20 hours. At each time point, digested peptides are removed from the system by a 3.5k MWCO dialysis membrane. The digested peptides at each time point are separated and identified by LC-MS/MS.

**Figure 3.**
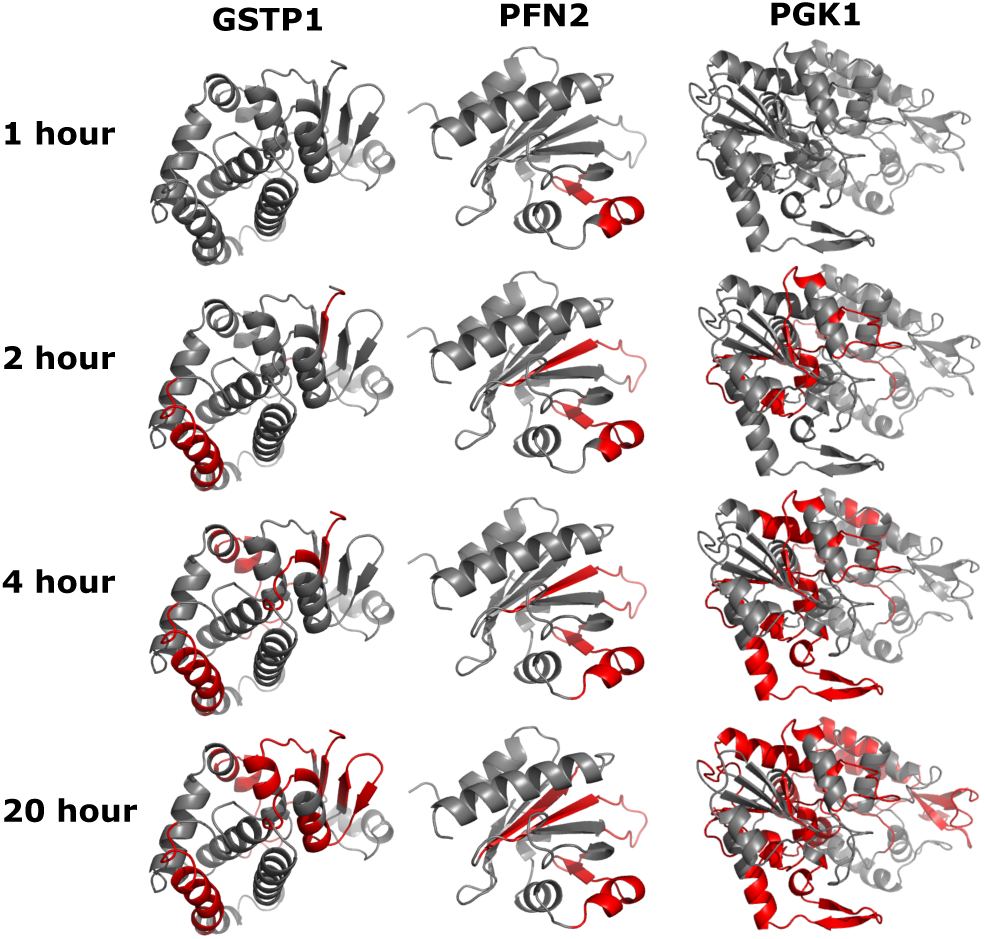
Limited proteolysis of native proteins visualized by SCV. Glutathione S-Transferase Pi 1 (GSTP1), Profilin 2 (PFN2), Phosphoglycerate Kinase 1 (PGK1) are selected as examples here. Red color highlights the detected peptides at each of the four time points, grey color shows the undetected portion of the protein.

When we combine the data from all four time points, and visualize them by SCV, we can easily see the progression of the digestion of trypsin (**Fig. 3**). In the three examples shown in **Fig. 3**, proteins are gradually digested from the outer surface to the inner regions. In the case of GSTP1, the outer regions are first observed as early as 2-hour. More inner regions of the GSTP1 are then followed in 4-hour and 20-hour time points. PFN2, on the other hand, is digested first at one of the helix regions at the surface, then followed by beta-sheet region inside. At the final 20-hour time point, the innermost beta-sheet region of PFN2 is eventually observed. In the case of PGK1, some outer regions are digested only in late time points, showing a possible resistance to digestion or possibly an early unfolding. The early unfolding might be a result of the digestion of key connection regions at an early time point. We need to point out that the undetected regions (grey) are mostly not due to resistance to digestion, but rather caused by low detectability of the corresponding peptides by mass spectrometry, either due to low ionization or non-ideal cleavage site arrangement. This visualization (**Fig. 3**) clearly shows the power of SCV to assist the structural investigation by proteomics experiments.

### Validate predicted structure

The predicted protein structures by Alphafold2 and RoseTTAFold are proven to be more accurate than the previously published algorithms. However, many challenging structures cannot be accurately predicted, e.g., intrinsically disordered proteins. Existing prediction models are trained using the currently available 3D structures from X-ray crystallography, NMR, and Cryo-EM experiments and therefore perform very well on these known structures as they are included in the training set. These predictions may also provide satisfactory results for many proteins with unknown structures but mainly composed of domains with known structures. There is no easy way to directly validate the predicted results for the unknown proteins with many unknown domains as these unknown domains have never been characterized by experiment.

On the other hand, all predicted structures must follow the same physical rules as the other known proteins. The accessibility of each region, defined by the predicted structure, must be consistent with the experimental observation. This gives us a unique opportunity to use our limited proteolysis results to validate and compare different prediction results. Here, we used data from our limited proteolysis experiments and mapped all observed peptides onto the structure predicted by Alphafold2 and RoseTTAFold. In order to make a comparison with the known PDB structure, here we chose the human phosphatidylethanolamine-binding protein 1 (PEBP1) as our test protein and used SCV to generate the 3D structure time series on Alphafold2, RoseTTAFold, and PDB structures (**Fig. 4**).

**Figure 4.**
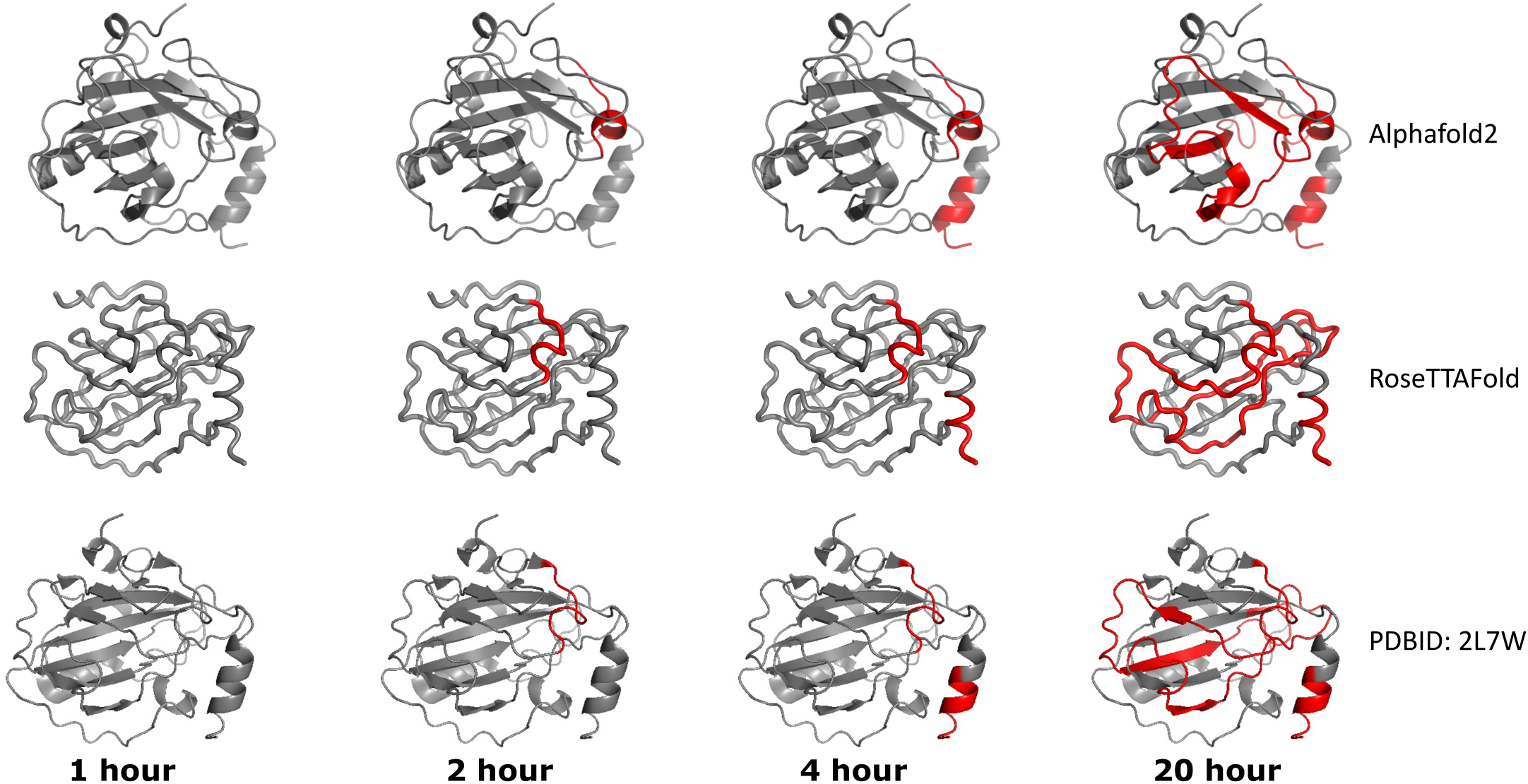
Comparison of Alphafold2, RoseTTAFold and PDB structure of phosphatidylethanolamine-binding protein 1 (PEBP1). Red color highlights the detected peptides at each of the four time points, grey color shows the undetected portion of the protein.

As shown in **Fig. 4**, both Alphafold2 and RoseTTAFold structures are quite similar to the PDB structure, with some differences. However, when we overlayed the limited proteolysis results on top of these structures, we can see that the differences between the two predicted structures are actually significant, especially in the upper left corner of the 20-hour digestion data. The 20-hour digestion data suggests that the RoseTTAFold predicted structure is more similar to the PDB entry. The Alphafold2 result with a slightly different angle is visually different than the other two. To validate this finding, we calculated the median distance change of all the identified residues from the 4-hour data point to the 20-hour data point and confirmed that the RoseTTAFold structure resulted in a trend similar to the PDB structure, while the Alphafold2 result is a bit different for this particular structure of PEBP1. The validation of unknown structure predictions is currently under investigation and will be published elsewhere.

In conclusion, SCV and limited proteolysis of native proteins can help us to visualize and compare different prediction results. We hope our data and the tool can help to validate and further improve protein structure prediction accuracy.

## Conclusion

Here we present a free, convenient, and powerful web application for the visualization of protein sequence coverage in 3D. We have shown that this tool is capable of visualizing PTMs, isotope labeling, and time-series data of limited proteolysis. The 3D visualization of limited proteolysis data may serve as an important step towards the validation of predicted structures from both Alphafold2 and RoseTTAFold. Overall, we demonstrated SCV could be easily used to obtain structural insights from proteomics results.

## Materials and Methods

All materials are purchased from Sigma-Aldrich unless otherwise mentioned.

### General material and cell

HEK 293T cells were cultured from frozen stock at 37°C in Dulbecco’s Modified Eagle Medium (high glucose DMEM, Sigma-Aldrich) supplemented with 10% Fetal Bovine Serum (FBS) and antibiotics (100 ug/ml penicillin and streptomycin). Cell passaging was conducted routinely twice a week when ∼80% confluency was reached. To prepare native protein lysate, ∼15 million cells were harvested and washed with 1* phosphate-buffered saline (PBS, pH 7.4). Sonication of the HEK cells diluted in 1* PBS was performed on ice at 30%, 60 kHz for 30 seconds with a repetition of 3 times and 15 seconds interval was given to cool down the lysate between each sonication. Native protein was precipitated by CHCl3/MeOH method after spin down at 13,000g at max speed, precipitated protein was air-dried and dissolved in 100mM Tris buffer (pH 7.4) and subsequential BCA assay (Invitrogen) was performed to measure native protein concentration prior to store at -80°C.

### Time-lapse limited proteolysis

To perform the limited proteolysis, 3 mg native HEK protein dissolved in 100mM Tris buffer was mixed with 150 ug trypsin (Promega, TPCK treated, 1:20), and 1mM CaCl2. The mixture was injected into a Slide-A-Lyzer dialysis cassette (3.5K MWCO, Thermo Scientific) using instructions provided by vendor. The dialysis cassette was then put directly into a transparent ziploc bag (generic, 9 cm by 5 cm) filled with 5 mL 100mM Tris buffer (pH 7.4) to ensure that after leftover air inside ziploc bag was squeezed out, dialysis cassette could be submerged by Tris buffer. The zipped ziploc was then incubated at 37°C for 1 hour, supernatants containing digested peptides from 1 hour were transferred to a clean tube and digestion continued with the addition of another 5mL fresh 100mM Tris buffer into the same ziploc bag. The supernatants corresponding to digested peptides from 2 hour, 4 hour and 18 hour were collected the consistent way as 1 hour sample. All collected supernatants were acidified upon time-point completion and desalted on C18 desalting columns (Thermo Scientific) following the manufacturer’s instructions and eluted in 50% acetonitrile (ACN) with 0.1% Trifluoroacetic acid (TFA). Peptides were speed-vac (vendor) at room temperature and reconstituted in 0.1% formic acid. Peptide concentrations were measured by a Nanodrop instrument (Thermo Scientific) at absorbance 205 nm.

### Proteomics data acquisition

A total amount of 2 μg equivalent of peptides from 4 time points were injected and separated by a capillary C18 reverse-phase column of the mass-spectrometer-linked Ultimate 3000 HPLC system (Thermo Fisher) using a 90-minutegradient (5-85% acetonitrile in water) at 300 nL/min flow. Mass spectrometry data were acquired on a Q Exactive HF mass spectrometry system (Thermo Fisher) with nanospray ESI source using data-dependent acquisition.

### Proteomics data analysis

Mass spectrometry raw files were converted into mzML files using MSconvert^10^, and searched by MSFragger^11^. The database used was a standard human reference proteome (20,371 reviewed entries only, UniProt ID: UP000005640, last modified on 3/7/2021) with 20,371 decoy sequences amending ^12^. Trypsin was specified as cleavage enzyme allowing up to 2 missing cleavage, mass tolerance was set to 50 ppm for precursor ions and 20 ppm for fragment ions. The search included variable modifications of methionine oxidation and N-terminal acetylation without any fixed modification. Peptide length was set within a range between 7 to 50 amino acids and false discovery rate (FDR) was set to 1% at the peptide spectrum match (PSM), peptide, and protein levels. In order to refine the predicted Alphafold2 structures, the depth of coverage at each time point was quantified since theoretically deeper coverage should be achieved with the prolonged digestion. Specifically, a custom Python script was used to map all identified peptides to sequence of protein of interest and coverage depth was computed as average Euclidean distance of all covered residues to the core of protein.

### 3D sequence coverage visualizer web application

Custom Python scripts were implemented to map all identified peptides including PTMs to proteome sequences, then residue 3D coordinates from PDB file, and residue positions of mapped region and PTMs were captured and outputted into a JSON object along with user-defined color representations processed with GLmol, which is dependent upon WebGL, a JavaScript API for rendering interactive 2D and 3D graphics within any compatible web browser.

A Node.js server accepts user input from an HTML form with a newline-separated list of PSMs, detected PTMs, and optional user uploaded PDB files as the fields. The PTMs are detected with a regular expression capturing the residue and mass within the sequence. The HTML form appended a generated universally unique identifier (UUID) in which the results may be accessed later. The Node.js server spawns the custom Python scripts with the fields of the user form as input to generate the resulting JSON object to be sent back to the user.

Client-side JavaScript renders a sorted list of identified proteins from the custom Python scripts by the percent sequence coverage. For each identified protein, a HTML canvas is instantiated and highlights the covered residues in red and PTMs in custom colors.

## Author Contributions and Acknowledgements

### Author Contributions and Notes

Y.G. and X.S. designed research, X.S. performed experiments, C.G. and X.S. wrote software, Y.G. and X.S. analyzed data; and Y.G and X.S. wrote the paper. The authors declare no conflict of interest.

## Acknowledgments

This work is supported by NIH grant 5R35GM133416 to Y.G.

